# simCAS: an embedding-based method for simulating single-cell chromatin accessibility sequencing data

**DOI:** 10.1101/2023.02.13.528281

**Authors:** Chen Li, Xiaoyang Chen, Shengquan Chen, Rui Jiang, Xuegong Zhang

**Affiliations:** MOE Key Laboratory of Bioinformatics and Bioinformatics Division of BNRIST, Department of Automation, Tsinghua University, Beijing 100084, China; School of Mathematical Sciences and LPMC, Nankai University, Tianjin 300071, China; Center for Synthetic and Systems Biology, School of Life Sciences and School of Medicine, Tsinghua University, Beijing 100084, China

## Abstract

Single-cell chromatin accessibility sequencing (scCAS) technology provides an epigenomic perspective to characterize gene regulatory mechanisms at single-cell resolution. With an increasing number of computational methods proposed for analyzing scCAS data, a powerful simulation framework is desirable for evaluation and validation of these methods. However, existing simulators generate synthetic data by sampling reads from real data or mimicking existing cell states, which is inadequate to provide credible ground-truth labels for method evaluation. We present simCAS, an embedding-based simulator, for generating high-fidelity scCAS data from both cell-wise and peak-wise embeddings. We demonstrate simCAS outperforms existing simulators in resembling real data and show that simCAS can generate cells of different states with user-defined cell populations and differentiation trajectories. Additionally, simCAS can simulate data from different batches and encode user-specified interactions of chromatin regions in the synthetic data, which provides ground-truth labels more than cell states. We systematically demonstrate that simCAS facilitates the benchmarking of four core tasks in downstream analysis: cell clustering, trajectory inference, data integration, and *cis*-regulatory interaction inference. We anticipate simCAS will be a reliable and flexible simulator for evaluating the ongoing computational methods applied on scCAS data.

**Availability:** simCAS is freely available at https://github.com/Chen-Li-17/simCAS.

## 1 Introduction

Rapid advances in single-cell sequencing technologies have enabled the characterization of cellular heterogeneity and identification of disease-specific processes at the single-cell level (Olsen and Baryawno, 2018). A range of single-cell chromatin accessibility sequencing (scCAS) technologies have been developed to study chromatin accessibility and gene regulation in single cells, mainly single-cell Assay of Transposase Accessible Chromatin with high-throughput sequencing (scATAC-seq) (Buenrostro, et al., 2015) and single-cell combinatorial indexing ATAC-seq (sci-ATAC-seq) (Cusanovich, et al., 2015). Specially, scATAC-seq can generate data from hundreds of thousands of cells on the timescale of weeks (Lareau, et al., 2019), which is an effective technology to dissect the activities of functional DNA sequences within specific tissues.

Multiple computational methods have been proved to be efficient in revealing the cellular heterogeneity in scCAS data (Fang, et al., 2021; Granja, et al., 2021), while benchmarking these methods quantitively with datasets of exact ground truths is still a tough challenge. For example, the unsupervised cell clustering methods utilize datasets with annotated cell types for evaluation (Kiselev, et al., 2019), while the cell types in real datasets are generally annotated manually and without external validation, which may bring unexpected artificial biases. Besides, taking the coarse-grained biological knowledge as ground truths may also lead to distortions in method benchmarking. For benchmarking the methods of reconstructing differentiation trajectories (Miao, et al., 2021) in scCAS data, the developmental relationships among different cell groups are provided as the ground truth, while these relationships are incapable of locating each cell on the developing trajectory. In addition to methods of identifying cell states, other analysis methods, such as data integration methods (Korsunsky, et al., 2019) and *cis*-regulatory inference methods (Li, et al., 2020; Pliner, et al., 2018) also require scCAS datasets with ground truth labels for better benchmarking. Therefore, a systematic and flexible simulator which provides synthetic data with exact and fine-grained ground truths for scCAS data will significantly facilitate the evaluations of analysis methods. However, due to the inherent high dimensionality of accessible peaks and sparsity of sequencing reads per cell (Chen, et al., 2019), the simulation for scCAS data remains a substantial challenge.

Compared to abundant simulation methods for scRNA-seq data (Cao, et al., 2021; Crowell, et al., 2022; Sun, et al., 2021; Zappia, et al., 2017), the existing several simulation methods for scCAS data are inadequate to satisfy the needs to benchmark diverse analyses. To our best knowledge, there are only four methods that can be used for simulating scCAS data: SCAN-ATAC-Sim (Chen, et al., 2021), simATAC (Navidi, et al., 2021), EpiAnno (Chen, et al., 2022) and scMultiSim (Li, et al., 2022). SCAN-ATAC-Sim generates scATAC-seq reads by taking bulk samples as input. Due to directly sampling the reads from bulk samples for each cell, SCAN-ATAC-Sim is unable to capture the characteristics of single cells. simATAC is the first simulator trained with single-cell data and generates synthetic data with discrete cell types of real data, while simATAC concentrates on modeling bins, the fixed chromatin regions, instead of peaks, the *cis*-regulatory elements with specific biological characteristics. EpiAnno solely simulates the peak-by-cell matrix with highly accessible peaks, and such a synthetic matrix without simulating full peak set is limited for various analyses. scMultiSim generates multi-modality data of single-cells with user-defined cell states, while the random selection of values in real scCAS data results in little resemblance of synthetic cells to real cells. In summary, none of existing methods provides a systematic simulation framework to generate a peak-by-cell matrix with user-defined cell states while maintaining the resemblance to real scCAS data.

To fill this gap, we propose simCAS, an embedding-based simulation framework that simulates scCAS data from low-dimensional embeddings with the user-defined settings. Our simulation framework can provide simulated scCAS data with unbiased ground-truth labels, such as cell states, data batches and *cis*-regulatory interactions. With the correction by the estimated statistics, simCAS generates data of superior resemblance to real data against existing simulators. To maintain the biological characteristics in *cis*-regulatory elements, simCAS provides a simulated peak-by-cell matrix with user-specific number of cells and number of peaks. By modulating the generation of cell-wise low-dimensional embeddings, simCAS generates data with ground-truth cell populations and differentiation trajectories, which significantly facilitates the benchmarking of analysis methods on identifying cell states. Moreover, the batch effects and *cis*-regulatory interactions can be optionally encoded in the synthetic data via adding Gaussian noise and moderating peak-wise embeddings generation, respectively, which extends the flexibility of our simulation framework. At last, we demonstrate the reliability and robustness of data generated by simCAS in benchmarking four computational tasks for scCAS data analysis: cell clustering, trajectory inference, data integration and *cis*-regulatory interaction inference.

## 2 Methods

### 2.1 simCAS framework

simCAS is an embedding-based method for scCAS data simulation (Fig. 1). To enable multi-scenario applications, simCAS provides three simulation modes, namely pseudo-cell-type mode, discrete mode and continuous mode, to generate synthetic data with pseudo-real manifold, discrete clusters and continuous differentiation trajectories, respectively. For the pseudo-cell-type mode, the input of simCAS is the real scCAS data represented by a peak-by-cell matrix, and matched cell type information represented by a vector. For the discrete or continuous mode, simCAS only requires the peak-by-cell matrix as the input data, followed by automatically obtaining the variation from multiple cell states. The output of simCAS is a synthetic peak-by-cell matrix with a vector of user-defined ground truths.

**Fig. 1.**
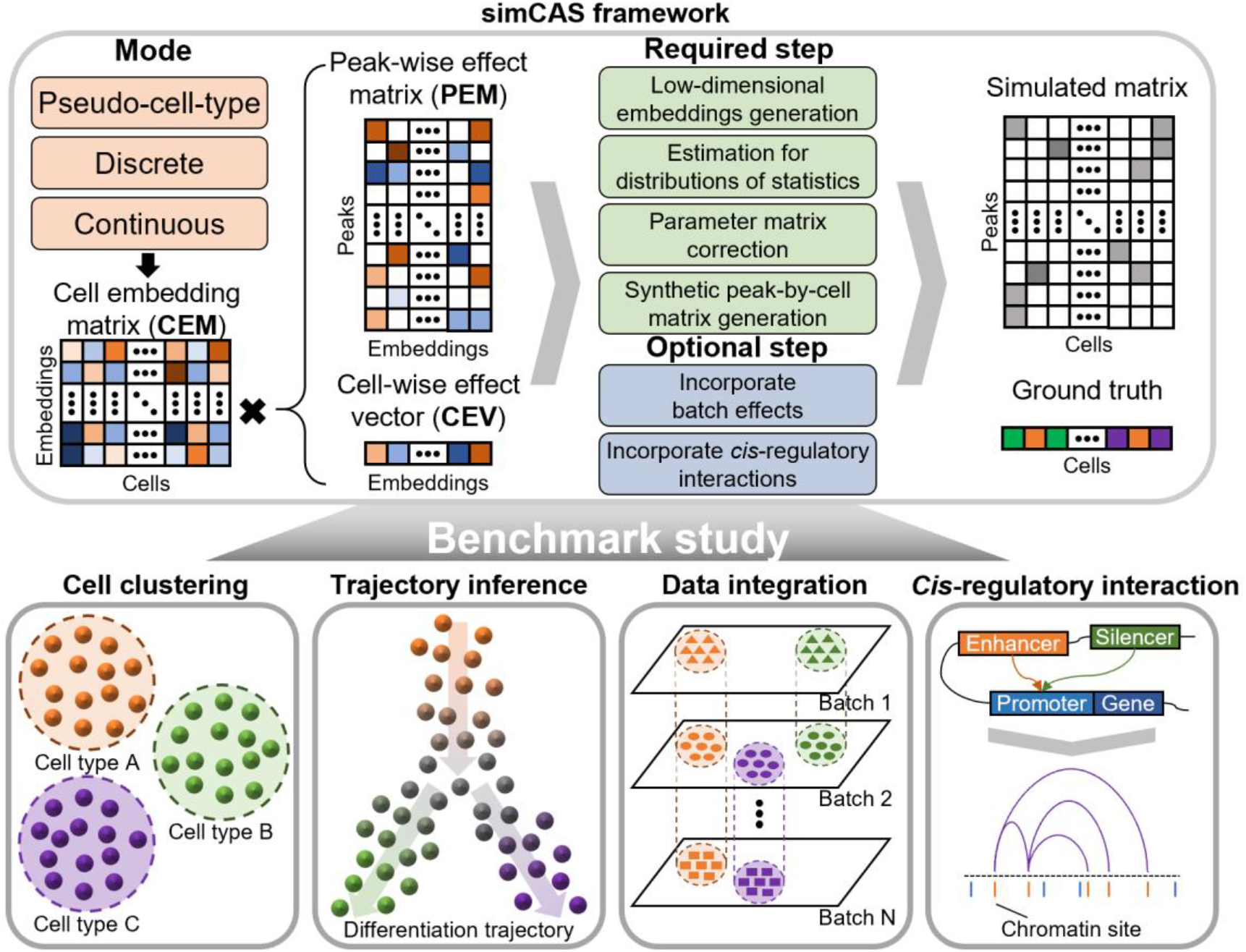
A graphical illustration of simCAS framework. simCAS provides three modes to simulate cells with different states: pseudo-cell-type mode, discrete mode and continuous mode. simCAS first generate cell-wise and peak-wise low-dimensional embeddings with user-defined ground truths. Then the distributions of statistics in real data are estimated for correcting the parameters. A peak-by-cell matrix is finally generated from the corrected parameters. The batch effects and interaction peaks can be added optionally during the simulation in different modes. The synthetic data generated by simCAS can be used to benchmark four major computational tasks of analyzing scCAS data: cell clustering, trajectory inference, data integration and cis-regulatory interaction inference.

For each mode, the required process of simCAS is consistent and can be divided into four major steps: (1) low-dimensional embeddings generation, specifically in cell-wise and peak-wise; (2) estimation for distributions of statistics, including the library sizes of the cells, the non-zero proportion of the cells, and the count summation of the peaks; (3) parameter matrix correction, by leveraging information from the aforementioned estimated distributions of statistics; (4) synthetic peak-by-cell matrix generation, that is, generating the final count matrix from a Poisson distribution. Note that a number of peak-by-cell count matrices are routinely binarized for downstream analyses, we thus provide an adapted framework with a Bernoulli assumption (supplementary Note 1). Batch effects and cis-regulatory interactions can be optionally added for task-specific benchmark studies.

#### 2.1.1 Low-dimensional embeddings generation

Given the number of simulated cells as *n*_*cell*_, the number of simulated peaks as *n*_*peak*_, and the dimension of cell embeddings as *n*_*embed*_, simCAS first generates two embedding matrices, namely cell embedding matrix (CEM) 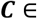 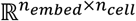 and a peak-wise effect matrix (PEM) 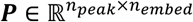, for characterizing the cell-wise and peak-wise features, respectively, and then generates a cell-wise effect vector (CEV) 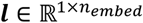, for determining the library size of each simulating cell. The CEM ***C*** can be divided into two sub-matrices, that is ***C*** = (***C***^*homo*^; ***C***^*hete*^), where ***C***^*homo*^ serves as the homogeneous CEM and ***C***^*hete*^ serves as the heterogeneous CEM. The homogeneous CEM ***C***^*homo*^ represents inherent cellular properties shared across all the cells, such as the locating tissue and the routine regulatory mechanism, while the heterogeneous CEM ***C***^*hete*^ crystallizes different cellular biological factors, such as differential chromatin accessibility and cell-specific developing state (Zhang, et al., 2019). The PEM ***P*** reflects the effects of cell embeddings to associated peaks. Elements in the CEV ***l*** control library sizes of simulated cells by setting different effect degree of each cell embedding.

The generation procedure of the homogeneous CEM ***C***^*homo*^, the PEM ***P*** and the CEV ***l*** is consistent among different modes. For the homogeneous CEM, we assume that each element in ***C***^*homo*^ follows a Gaussian distribution with a unit mean and a user-defined variance *σ*^2^, which determines the extent of data dispersion. As a side note, the parameter of variance *σ*^2^ is consistent between the homogeneous CEM and the heterogeneous CEM. For generation of the PEM, simCAS draws each element in ***P*** from a standard Normal distribution, followed by randomly setting the elements to zero with probability *η* by row. *η* is set to 0.5 in this study. For generation of the CEV, values of elements in ***l*** are randomly sampled from a standard Gaussian distribution.

For the heterogeneous CEM, we denote 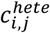 as the element of the *i*th embedding and *j*th cell in 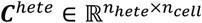, where *n*_*hete*_ is the dimension number of heterogeneous embeddings. The generation of heterogeneous CEM determines the cell states to be discrete or continuous in the final synthetic matrix. For the discrete mode, *n*_*pop*_, that is the number of discrete cell populations, and a covariance matrix 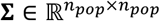, accounting for the distance between different cell populations, are needed as user input. Then *n*_*pop*_ vectors with *n*_*embed*_ dimensions are sampled from multivariate normal distribution 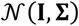 and concatenated to a matrix 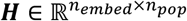, and the heterogeneous CEM value of *i* th embedding and *j*th cell within cell population *k* is generated by:

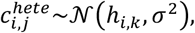

where *h*_·,·_ is the element in ***H***. For the continuous mode, simCAS requires users to input a Newick tree-format data with tree nodes and branch lengths. simCAS first assigns the cells to each branch in proportion to the branch length and set a uniform interval for adjacent cells on the same branch. Then simCAS generates 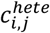 with a Brownian motion assumption:

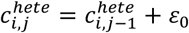

where (*j* − 1)th cell is the previous cell of *j*th cell from root to leaves along the tree, and *ε*_0_ is an increment sampled from a normal distribution with a zero mean and the variance equal to the interval between *j*th cell and (*j* − 1)th cell. For the root cell, the value of 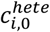 is sampled from a normal distribution with a unit mean and the aforementioned variance *σ*^2^. It is worth mentioning that there is no heterogeneous part in CEM for the pseudo-cell-type mode, because the heterogeneous information has been included in the input data. In this study, *n*_*embed*_ and *n*_*hete*_ are fixed to 12 and 10, respectively.

#### 2.1.2 Estimation for distributions of statistics

Given a real peak-by-cell count matrix, simCAS estimates distributions of three core statistics from real data: library size (the number of aligned reads per cell), peak summation (the sum of aligned reads per peak), and cell non-zero proportion (the proportion of non-zero values per cell).

For the pseudo-cell-type mode, similar to simATAC (Navidi, et al., 2021), simCAS models log-transformed library size and log-transformed cell non-zero proportion by two Gaussian mixture models (GMM) with two components, respectively. To estimate the distribution of peak summation, simCAS fit a variant of Logarithmic distribution (referred to as Log-variant distribution), which has a characteristic of long right tail as with the real (Fisher, et al., 1943). For simplicity of notation, we assume *x* as a random variable with no practical meaning in the following discussion. With specific modeling for the count of zero and one, the probability mass function of the Log-variant distribution is:

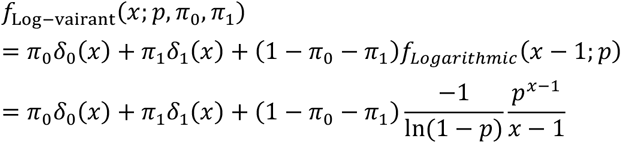

where *δ*_0_(·) and *δ*_1_(·) indicate the point mass at zero and one, respectively, and *π*_0_ and *π*_1_ are the corresponding probabilities. *p* is a parameter ranging from 0 to 1 in Logarithmic distribution. We also provide other five discrete distributions for modeling peak summation as alternatives (Supplementary Note 2).

For the discrete or continuous mode, simCAS uses kernel density estimation (KDE) to estimate the distributions of statistics (Zhang, et al., 2019) so that it is adaptive to the diversity of input mixing cell types.

#### 2.1.3 Parameter matrix correction and synthetic peak-by-cell matrix generation

By multiplying the PEM 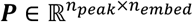 and the CEM 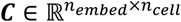, we will obtain a parameter matrix 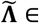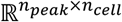 with the same shape as the final output matrix. Note that there is no extra restriction of PEM and CEM except for sampling from Gaussian distributions, part of elements in 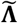 are inevitably negative and cannot be directly used as the parameter in Poisson distributions. To transform 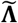 into an expected parameter matrix, simCAS performs the two following operations: (1) transform elements in 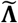 into positive via an activation function; (2) perform cell-wise and peak-wise correction using the fitted distributions in the previous step.

First, we developed two activation functions adaptive to different modes. For the discrete or continuous mode, we provide a piecewise function *f*_1_(·) by integrating an exponential function and a linear function as the activation function:

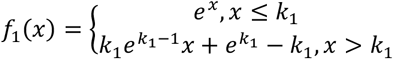

where *k*_1_ is a parameter to control the variance of simulated data, and a higher *k*_1_ value brings higher variance of peak accessibility. In this study *k*_1_ is fixed to 2. For pseudo-cell-type mode, we expect the diversity within a cell type less than between different cell states, and a Sigmod-format function *f*_2_(·) with a smaller slope is provided as the activation function:

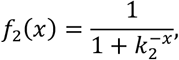

where *k*_2_ is a parameter to adjust the steepness of the activation function curve and fixed to 2 in this study. After this operation, the parameter matrix 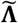 is transformed into an activated parameter matrix 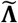.

Second, simCAS conducts correction with estimated statistical distributions of library size, peak summation and cell non-zero proportion in turn, to make simulated data preserve the cell-wise and peak-wise properties as with the real. For instance, simCAS performs the library size correction as follows:

Multiply the CEV ***l*** and the CEM ***C***, and obtain 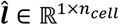.

Randomly sample *n*_*cell*_ values from the GMM distribution fitted with real library sizes to form the synthetic library size set 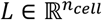.

Sort the elements of 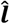 and values in *L* to obtain the sorting indices.

Obtain the vector 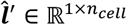 of the library sizes of synthetic cells by replacing the elements in 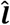 with the sampled values from *L* one by one according to the sorting indices.

For each column vector (represents each cell) in 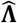, divide each element by the sum of the vector, multiply all elements by the corresponding element (represents the associated cell) in 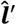, and obtain the corrected parameter matrix 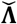.

Details of peak summation and cell non-zero proportion corrections are provided in Supplementary Note 3 following a similar correction idea. By the two operations, simCAS obtains the mean parameter matrix **Λ**, of which elements are served as mean parameters of Poisson distributions.

In the synthetic peak-by-cell matrix generation step, each element of the final simulated count matrix 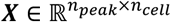 is drived from a Poisson distribution one by one with the corresponding mean parameter in **Λ**.

#### 2.1.4 Optional steps in simCAS

To facilitate benchmark studies of various downstream analysis, simCAS offers several optional steps, more specifically, to incorporate batch effects or cis-regulatory interactions with synthetic data in the step of low-dimensional embeddings generation. Batch effects, namely unwanted variations in single-cell sequencing data of various batches, will potentially interfere with the biological analysis. The effects can be divided into two major categories (Luecken, et al., 2022): technical variations, mainly in sample composition, sequencing technologies and more; biological factors, such as spatial locations, tissues and species. In simCAS, users could add and adjust Gaussian noises to the mean parameter matrix **Λ** and the PEM ***P***, respectively, for simulating batch effects of technical variations or biological factors. Furthermore, simCAS can also generate data with user-defined cis-regulatory interactions by remodeling the PEM. Details of the optional steps in the simCAS framework are provided in Supplementary Note 4.

### 2.2 Data collection and pre-processing

We follow the standard pipeline of epiScanpy (Danese, et al., 2021) to filter out peaks that covered in too few cells and cells that do not have enough accessible peaks, and then remove cell types with less than 50 cells. We collected four scCAS datasets with peak-by-cell matrices and matched cell-type labels: Buenrostro2018 (Buenrostro, et al., 2018), Li2021 (Li, et al., 2021), Preissl2018 (Preissl, et al., 2018) and Chiou2021 (Chiou, et al., 2021). A summary of the above datasets after preprocessing is shown in Table 1.

**Table 1.** Summary of datasets used in this study.

| Dataset | Tissue | Protocol | Number of cells | Number of peaks | Number of cell types | Data accession |
| --- | --- | --- | --- | --- | --- | --- |
| Buenrostro2018 | Blood (human) | scATAC-seq (Fluidigm C1) | 1,931 | 169,221 | 10 | GSE96772 |
| Li2021 | Cortex (mouse) | snATAC-seq | 5,532 | 154,308 | 16 | GSM5273008 |
| Preissl2018 | Forebrain (mouse) | snATAC-seq | 2,062 | 115,763 | 10 | GSE100033 |
| Chiou2021 | Islet (human) | snATAC-seq | 15,298 | 91,754 | 12 | GSE160472 |

### 2.3 Other methods used in this study

For baseline methods, we implemented two simulators with source code obtained from their studies: simATAC (Navidi, et al., 2021), the first scATAC-seq simulator as we known, and scMultiSim (Li, et al., 2022), a simulator for single cell multi-omics generation. Since both simATAC and scMultiSim can only simulate data with the same manifold as real, we compare the performance of these methods with simCAS in the pseudo-cell-type mode. Note that EpiAnno (Chen, et al., 2022) only generates data with highly accessible peaks instead of whole peaks, and SCAN-ATAC-sim (Chen, et al., 2021) requires bulk data as input, so we exclude these two simulators as baseline methods.

For benchmarking clustering methods, we tested Leiden clustering, k-means clustering and Hierarchical clustering (HC) methods (Chen, et al., 2019) using the simulated data generated by simCAS in the discrete mode. For benchmarking trajectory inference methods, we tested Monocle3 (Cao, et al., 2019) with different parameters. For benchmarking methods of data integration or cis-regulatory interaction inference, we also tested Harmony (Korsunsky, et al., 2019) and Cicero (Pliner, et al., 2018) on synthetic data with multiple batches and cis-regulatory interactions, respectively.

For data visualization, we first select 50000 highly accessible peaks, then perform term frequency-inverse document frequency (TF-IDF) and principal component analysis (PCA) transformation to reduce dimensions to 50, and finally apply uniform manifold approximation and projection (UMAP) to project the cells into a 2-dimensional space. Unless otherwise stated, we perform the above pipeline for visualization in this study.

### 2.4 Metrics for evaluation

We assess the simulation performance from two perspectives, namely statistical evaluation and biological evaluation. For statistical evaluation, we focus on the average of read counts per peak (peak mean), the library size and the zero-peak proportion in each cell (cell sparsity) as with simATAC, and measure the diversity between the simulated data and real data by calculating median absolute deviation (MAD), mean absolute error (MAE), root mean square error (RMSE), Pearson correlation coefficient (PCC), Jensen-Shannon divergence (JSD), Kolmogorov-Smirnov statistic (KSS). As a side note, for the library size, we perform log-transformation before comparison as with the recent benchmark studies (Cao, et al., 2021; Crowell, et al., 2022). For biological evaluation, we compute median integration local inverse Simpson’s index (miLISI) (Sun, et al., 2021) to quantify the similarity between synthetic cells and real cells.

To evaluate the performance of different clustering methods on synthetic data generated by simCAS, we perform the following three metrics: adjusted mutual information (AMI), Homogeneity score (Homo) and adjusted Rand index (ARI) (Chen, et al., 2019). To quantitively benchmark the performance of cis-regulatory interaction inference methods, we take interactive peaks as positive samples and non-interactive peaks as negative samples in a selected peak hub, and F1 score are used to assess the accuracy of annotation. Details of the above evaluation are provided in Supplementary Note 5.

## 3 Results

### 3.1 simCAS generates high-fidelity cells with consistent manifold with real data

To demonstrate the advantage of simCAS with the pseudo-cell-type mode for data simulation, we conducted the statistical evaluation and biological evaluation using four datasets (Methods). In this section, simCAS was benchmarked against two baseline methods, simATAC and scMultiSim. Using the peak-by-cell matrix and the cell-type labels of each dataset as input, simCAS and baseline methods simulated the peak-by-cell matrix for each cell type with the same number of peaks and cells as in the real datasets.

For statistical evaluation, we first perform comparisons of three properties, namely peak mean, library size, and cell sparsity of synthetic data to real by cell type (Methods). Fig. 2a and Supplementary Fig. 1 depict the comparison of the three statistics’ distributions in all cell types between the synthetic dataset and the real dataset (Buenrostro2018), and demonstrate that data generated by simCAS highly resemble real data at peak-wise and cell-wise. It is observed that when trained with more cells, sinCAS can better preserve these properties, indicating that simCAS better capture the cell-wise and gene-wise characteristics if provided with a larger cell population. To quantitively measure the similarity of statistics’ distributions between synthetic data and real data, we further calculated MAD, MAE, RMSE, 1-PCC, JSD and KSS (Methods). A smaller value of each metric means that the simulated peak-by-cell matrices more accurately preserve properties of real data. As shown in Fig. 2b, simCAS significantly outperformed simATAC and scMultiSim across all cell types in Buenrostro2018 dataset. Taking MAD as an example, for peak mean, library size, and cell sparsity, the average value of simCAS is 54.5%, 20.2% and 93.8% lower than simATAC, respectively, and 60.0%, 50.3% and 84.4% lower than scMultisim, respectively. Focusing on peak mean and library size, scMultiSim provided overall the worst performance with the highest diversity between synthetic data and real data. Due to neglecting to model the cell sparsity in scCAS data, simATAC leads to the most spurious estimation in this property. This is consistent with the observation in the study of EpiAnno that simATAC generated synthetic data as pseudo-bulk data instead of single-cell data (Chen, et al., 2022). On the other datasets, simCAS still showed the superiority in the statistical evaluation (Supplementary Fig. 2a, 3a and 4a). The details of the quantitive evaluation for all datasets are provided in Supplementary Table. By replacing the distribution for modeling peak summation with other discrete distributions, we demonstrate that Log-variant distribution achieves best performance for fitness overall in majority cell types (Supplementary Fig. 5). In addition to statistical evaluation at cell-wise and peak-wise, we also tested whether simCAS can capture the property of peak-peak correlations. We first selected top 2,000 peaks with highest accessibility in real dataset and preserved the same peaks in simulated datasets as above. We next calculated the Spearman correlation coefficients of every peak-peak pairs’ chromatin accessibilities in the processed real and simulated datasets. As shown in Fig. 2c, simCAS best captures the correlations among highly accessible peaks, while a higher value and a lower value of Spearman correlation coefficients are presented by simATAC and scMultiSim, respectively. We also used the other four datasets to show the performance of simCAS to robustly capture the peak correlations (Supplementary Fig. 2b, 3b and 4b).

**Fig. 2.**
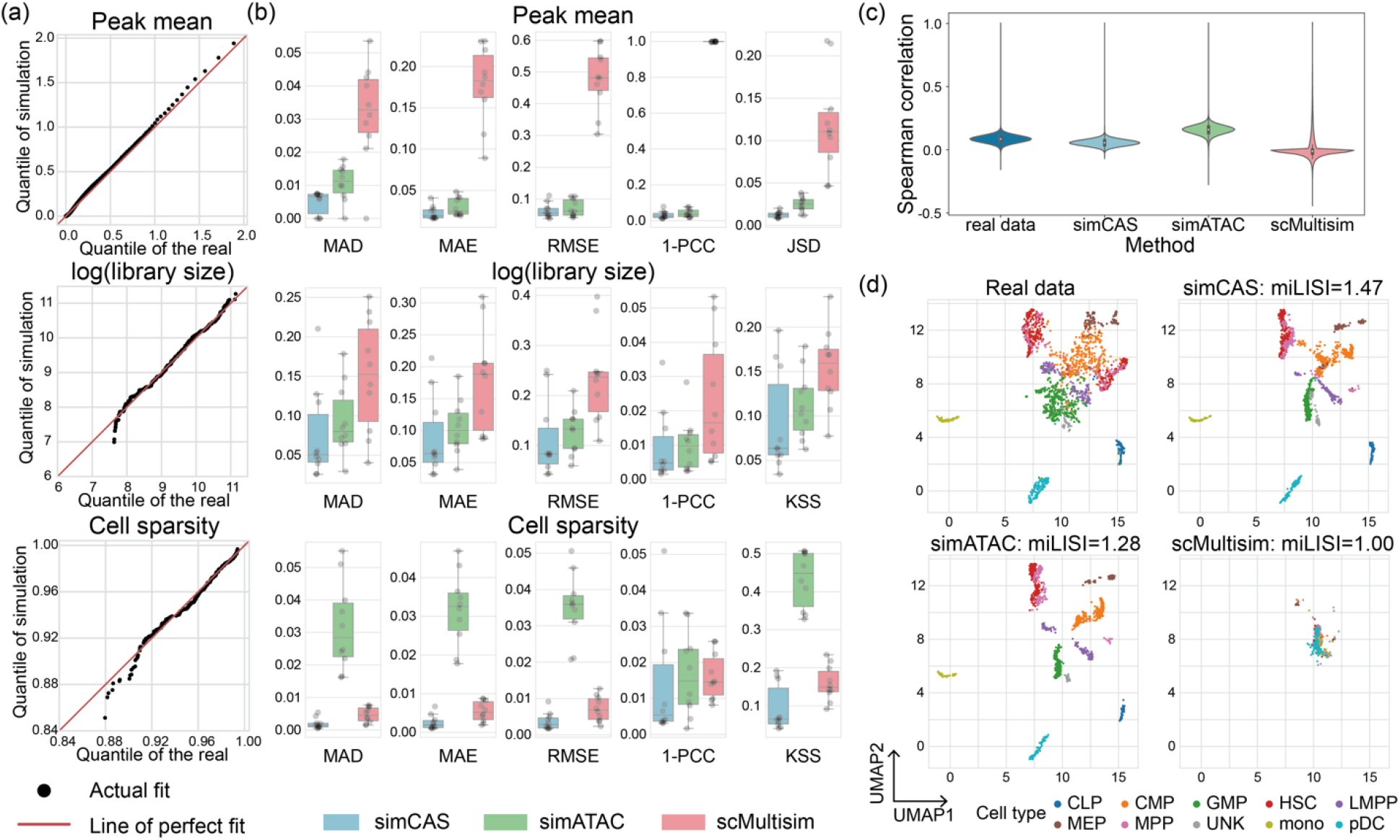
Comparisons of synthetic data and real data. (a) QQ-plots of peak mean values, library size values and cell sparsity values between synthetic common myeloid progenitors (CMPs) generated by simCAS and real CMPs in Buenrostro2018 dataset. 1000 quantiles of peak mean values are used for the comparison. (b) Comparison results of three statistics between synthetic cell types and real cell types in Buenrostro2018 dataset. For each cell type, the similarity between statistics of synthetic data and real data is measured by six metrics: MAD, MAE, RMSE, 1-PCC, JSS and KSS, and each gray point represents a cell type. (c) UMAP visualization of the synthetic datasets and real Buenrostro2018 dataset. The synthetic datasets are projected to the embedding space with the same low-dimensional projection trained on the real data. miLISI values are calculated to measure the mixture of synthetic cells and real cells for different simulators. (d) Spearman correlation coefficients of synthetic and real datasets on the pairs of top 2000 highly variable peaks selected in real data.

For biological evaluation, we assessed the similarity between simulated cells and real cells for each dataset by calculating the miLISI value of mixed cells (Methods). The results of data visualization and biological evaluation are shown in Fig. 2d, simCAS achieves highest miLISI value and, consequently, has superior simulation capability to capture the structure of cell clusters (Method). simATAC generates cells that gather to a small group within a certain cell type, which indicates that these simulated cells are less heterogeneous compared to the real cells. scMulitisim generates data without considering the diversity of different cell types and the visualizing result is a concentrated dense population in the UMAP space. The visualizations and miLISI values of other four dataset (Supplementary Fig. 2c, 3c and 4c) showed the satisfactory performance of simCAS to maintain the realistic manifold of scCAS data.

### 3.2 simCAS simulates data with discrete cell states to benchmark cell clustering methods

The cell types in most real scCAS data are obtained by unsupervised cell clustering and manual annotation, which highly rely on the investigator’s background knowledge, and may be not accurate enough as the groud truths for quantitive evaluation to analysis methods (Chen, et al., 2021; Chen, et al., 2022). Therefore, synthetic data with ground-truth labels of cell types will greatly benefit the increasing computational methods for cell clustering in scCAS data. We thus developed the discrete mode in simCAS for simulating cells with user-defined cell populations. Using the peak-by-cell matrix of Buenrostro2018 dataset as training data, we generated three synthetic datasets with different parameters: A1 dataset with the covariance matrix of different cell populations **Σ** = **Σ**_1_ and the standard deviation in CEM generation *σ* = 0.5, A2 dataset with **Σ** = **Σ**_2_ and *σ* = 0.5, and A3 dataset with **Σ** = **Σ**_2_ and *σ* = 0.7 (Fig .3a). We set the number of cells, the number of peaks and the number of populations to 1,500 (300 for each cell population), 169,221 (same as the number of real peaks) and 5, respectively, and keep these parameters in the three synthetic datasets. As the UMAP visualization in Fig. 3a, the inter-population distances among different populations maintain the relationships encoded in **Σ**_1_. When rising the covariance between population A and B from 0.56 in **Σ**_1_ to 0.78 in **Σ**_2_, cells between population A and B exhibit a closer relationship, and cells of population C are separated slightly from cells of population A and B. Besides setting the covariance matrix, some other parameters can be set to control the properties of simulated data. For example, a higher value of *σ* brings higher inner-population variance as shown in A2 and A3 dataset (Fig. 3a). To further demonstrate the high resemblance of synthetic data to real data in the discrete mode, we also showed that simCAS successfully retain the properties of peak mean, library size and cell sparsity (Fig. 3b and Supplementary Fig. 6a).

**Fig. 3.**
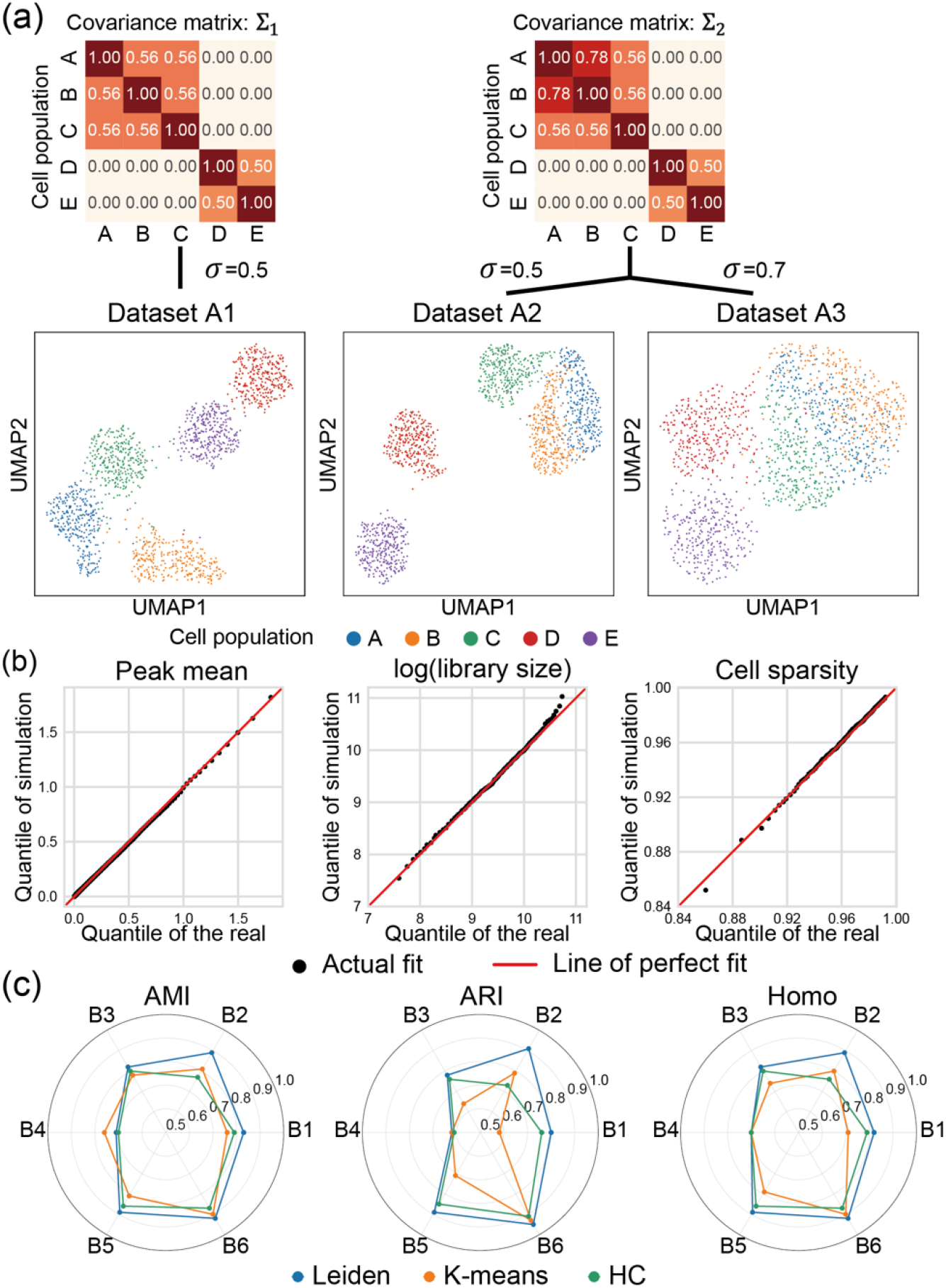
Synthetic data generation in discrete mode of simCAS and benchmarking of cell clustering methods. (a) UMAP visualization of synthetic datasets (A1-A3) with discrete cell populations generated by simCAS, colored by user-defined populations in the input covariance matrices. Dataset A1 and A2 are generated with different input covariance matrices. Dataset A2 and A3 are generated with different values of variance parameter. (b) QQ-plots of peak mean values, library size values and cell sparsity values between A1 dataset and real Buenrostro2018 dataset. Peak mean is compared using 1000 quantiles, and library size is compared using 100 quantiles as well as cell sparsity. (c) Performance benchmarking of Leiden clustering, K-means clustering and hierarchical clustering by AMI, ARI and Homo. Six simulated datasets (B1-B6) with discrete cell populations of cells number ranging from 500 to 3000 and populations number ranging from 3-7 are utilized for the benchmarking.

We then simulated 6 datasets (referred to as the B1 to B6 dataset) of discrete data with different number of cells (500 to 3000) and different number of populations (3 to 7) to evaluate three commonly used unsupervised clustering methods for single-cell analysis: Leiden clustering, K-means clustering and HC (Methods and Supplementary Fig. 7a). For Leiden clustering, we implemented a binary search to tune the resolution to match the number of populations and the number of clusters, while K-means clustering and hierarchical clustering are directly set the number of clusters to the number of cell populations. As shown in Fig. 3c, Leiden clustering and hierarchical clustering work consistently across different metrics, and Leiden clustering performs better than K-means clustering and hierarchical clustering in almost all the datasets. Taking B2 dataset as an example, we showed clustering results with the UMAP visualization in Supplementary Fig. 7b. The results of benchmarking analysis on our simulated data are consistent with the benchmark study for clustering methods on single-cell data (Chen, et al., 2019), indicating that data generated by simCAS is credible to benchmark the clustering methods on scCAS data with different sizes and dimensions.

### 3.3 simCAS simulates data with continuous differentiation trajectories

Although a range of methods have been developed to distinguish different cell types in scCAS data, the cellular processes are dynamic in nature and not always well described by these methods (Chen, et al., 2019). Therefore, methods for trajectory inference are explored to provide more comprehensive analyses of single-cell data (Saelens, et al., 2019). However, the information of cell trajectory relies on existing biological knowledge, which is imprecise and inconvenient to evaluate analysis methods for data with continuous trajectories. To fill this gap, simCAS provides the continuous mode to generate data by defining the cell trajectory with an input Newick tree. Using the Buenrostro2018 dataset as training data, we simulated three datasets as follows: C1 dataset with the Newick tree **T**_1_ and the standard deviation of CEM *σ* = 0.5, C2 and C3 datasets with the same tree **T**_2_ but with different standard deviations *σ* = 0.5 and *σ* = 1.0, respectively (Fig. 4a). For each dataset, we generated a peak-by-cell matrix with the shape of 169,221 and 1500. The UMAP visualization showed that, cells generated by simCAS explicitly maintains the trajectory structure from the input tree. More specifically, the length of trajectory in the UMAP space is in direct proportion of the branch in the input tree, and for example, cells of a shorter branch, such as ‘R-E’ in **T**_2_, are grouped to a smaller cluster. The value of *σ* also controls the dispersion of cells. We further compared the distributions of peak mean, library size and cell sparsity between simulated data and real data. As shown in Fig. 4b and Supplementary Fig. 6b, data generated by simCAS in the continuous mode highly resemble real data in cell-wise and peak-wise properties.

**Fig. 4.**
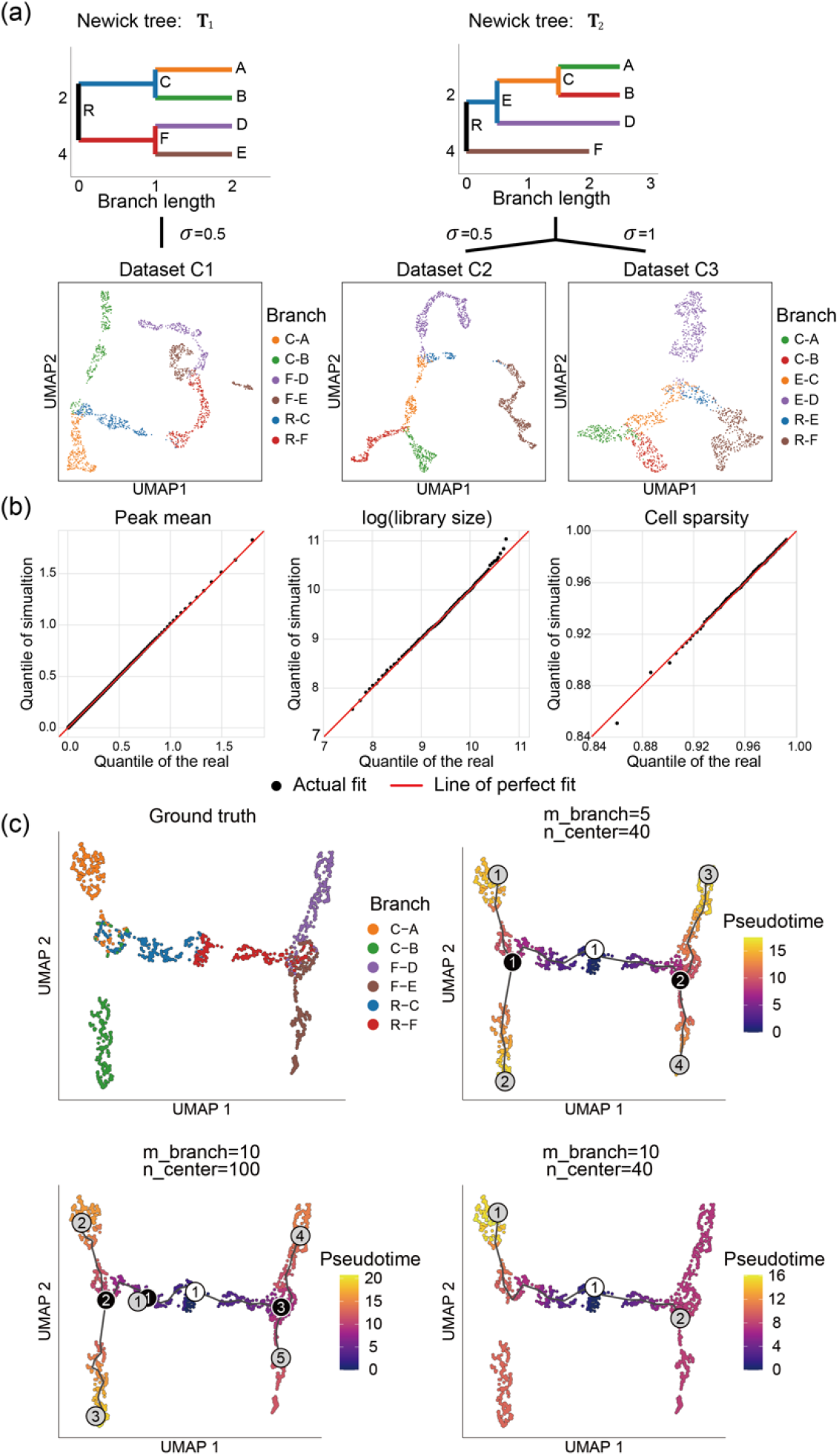
Synthetic data generation in continuous mode of simCAS and benchmarking of trajectory inference method Monocle3. (a) UMAP visualization of synthetic datasets (C1-C3) with continuous trajectories generated by simCAS, colored by the branches of user-defined input trees. Dataset C1 and C2 are generated with different input trees of specific nodes and branch lengths. Dataset C2 and C3 are generated with different values of variance parameter. (b) QQ-plots of peak mean values, library size values and cell sparsity values between C1 dataset and real Buenrostro2018 dataset. Peak mean is compared using 1000 quantiles, and library size is compared using 100 quantiles as well as cell sparsity. (c) Results of trajectory inference by Monocle3 applied on dataset C1. The performance of Monocle3 is evaluated with different values of two parameters, minimal branch length and number of centers. The inference results of cells are shown in the UMAP space, colored by the estimated pseudotime along with the inferred trajectories. White nodes, black nodes and gray nodes represent root nodes, branch nodes and leave nodes, respectively.

Using C1 dataset as input, we evaluated the parameter configuration in Monocle3, a method for trajectory inference and pseudotime estimation, for testing two fundamental parameters, minimal branch length (m_branch) and number of centers (n_center). We then used Monocle3 to visualize cells characterized by ground truths and predict trajectories with different parameters (Fig. 4c). The results indicated that a lower value of minimal branch length brings more branches of the trajectory structure, and a larger number of centers makes the trajectory structure more complicated, which are consistent with instructions in the study of Monocle3. Compared with other parameters, Monocle3 with minimal branch length of 5 and centers number of 40 completely delineate the real trajectory and detect the tree nodes. Altogether, simCAS provides a new perspective to computational method development for trajectory inference.

### 3.4 simCAS contributes to benchmarking in single-cell data integration

With the data integration methods applied on scCAS data, datasets from various origins can be analyzed simultaneously, which provides a comprehensive perspective to study cellular heterogeneity (Kopp, et al., 2022; Yuan and Kelley, 2022). Whereas, due to the challenges for distinguishing batch effects from indicative biological variances (Luecken, et al., 2022), it can be difficult to evaluate the methods of batch effect correction or data integration objectively. simCAS provides an opportunity to benchmark methods for this task by generating data with simulated batches. simCAS with the discrete mode can incorporate user-defined batch effects of biological factors, namely biological batch effects, with simulated data by adding Gaussian noise on the PEM (Methods). For instance, we first set the number of populations to 3 (the number of cells in cell population A, B and C is set to 600, 600 and 300, respectively) and a unit diagonal matrix as the covariance matrix, included a Gaussian noise with the mean of 0.5 and the standard variation of 0.5 to the PEM, and performed the remaining steps to generate a peak-by-cell matrix, regarded as Batch 1. We then performed the same procedure without adding Gaussian noise to generate another peak-by-cell matrix, regarded as Batch 2. Note that these two matrices are generated with exponential function to increase difference between the batches. Finally, we concatenated the two matrices by column, and obtained a synthetic dataset D1 with two batches. With the data visualization shown in Fig. 5a, we demonstrate that the simCAS effectively simulates biological batch effects within each cell population. Otherwise, to mimic batch effects of technical variations, that is technical batch effects, simCAS adds Gaussian noise the mean parameter matrix **Λ**, and the UMAP visualization illustrates the well-defined technical batch effects compared with cell populations (Supplementary Fig. 8).

**Fig. 5.**
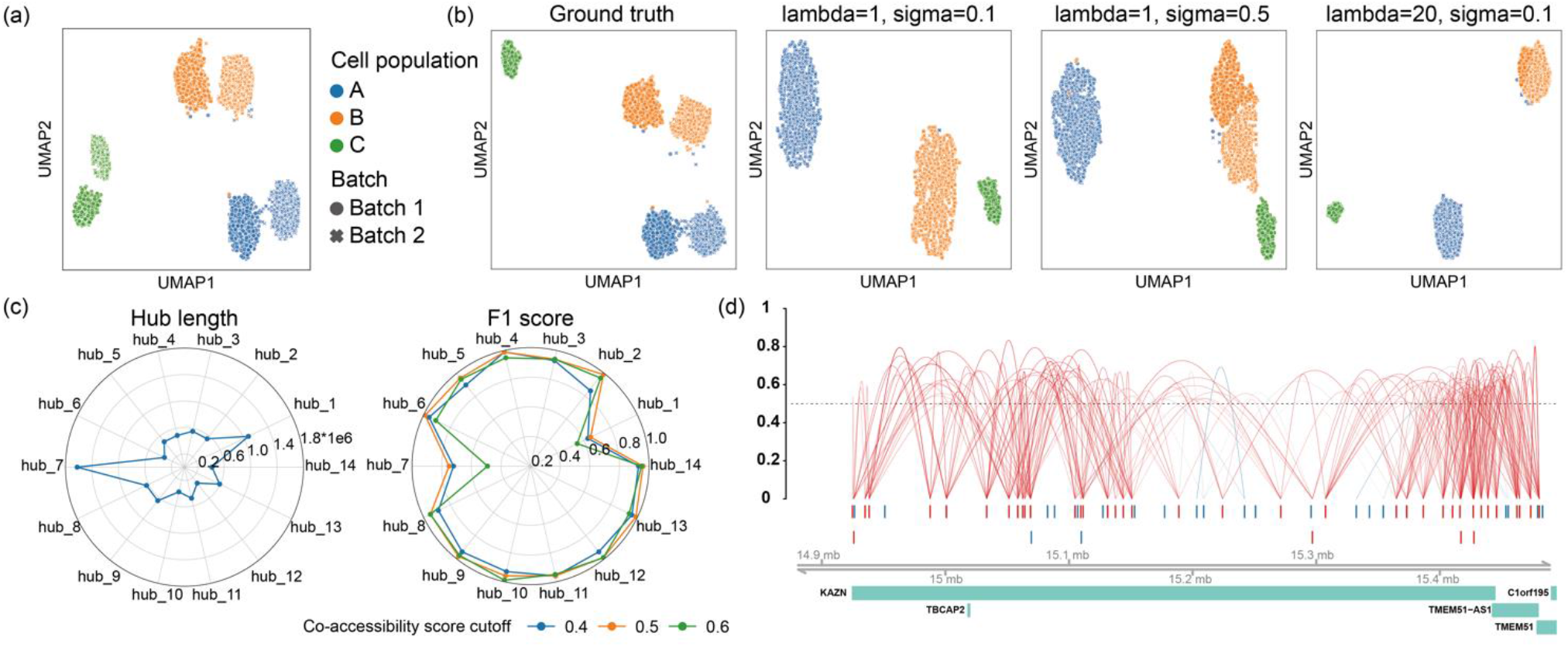
Method benchmarking results of data integration and cis-regulatory interaction inference. (a) UMAP visualization of synthetic dataset of 3000 cells with simulated biological batch effect, colored by the cell populations and marked by different batches. (b) Results of different parameter configurations of lambda and sigma in Harmony integration method. The integration results are visualized in the UMAP space, colored by cell populations and marked by simulated batches. (c) Length of selected 14 peak hubs, which represent high accessible gene regions, ranging from 337,196 bp to 1,622,989 bp (top). Cicero is applied to infer the interactive peaks defined in these hubs with co-accessibility score of 0.4, 0.5 and 0.6, and the performance is benchmarked with the F1 score calculated between predicted peaks in CCANs and ground truth interactive peaks in synthetic data (bottom). (d) The predicted connections by Cicero on the peak hub of gene *KAZN* region extended upstream by 50kb and downstream by 50kb. The 40 red peaks represent the ground truth interactive peaks defined in the synthetic data, and the remaining 26 blue peaks are non-interactive peaks.

To benchmark the parameter configurations of Harmony, a widely used integration method for single-cell sequencing data, we adapted D1 dataset to a more challenging dataset D2 by removing cell population C in batch2, to mimic the scenario with batch-specific and rare cell types. Two parameters of Harmony, namely lambda, a ridge regression penalty parameter, and sigma, the width of soft K-means clusters, are evaluated. As shown in Fig. 5b, the results of UMAP visualization demonstrate that, a larger value of sigma assigns cells to more clusters, while a smaller value of lambda results in more aggressive integration, as with the instruction in Harmony. With the lambda of 1 and sigma of 0.1, Harmony successfully integrates cells from different batches and separates cells of the different cell populations. With a larger sigma of 0.5, Harmony clusters cells to 4 groups and failed to correct the batch effects of cell population B, and with a larger lambda of 20, Harmony still separated different cell populations well, but the cells in population B are not well mixed between different batches. The parameter configuration evaluation of Harmony confirms the capability of simCAS to be a reliable simulator to benchmark data integration methods for scCAS data.

### 3.5 simCAS facilitates benchmarking in cis-regulatory interaction inference

scCAS data provides a perspective to investigate the interactions of chromatin accessible sites, and several computational methods (Dong and Zhang, 2021; Li, et al., 2020), such as Cicero (Pliner, et al., 2018), have been proposed to predict the interactions between peaks, namely cis-regulatory interactions. However, in the absence of fine-grained labels which contain correlations among all the chromatin sites, quantitative evaluation of the corresponding methods presents a unique challenge. We here conducted a series of experiments to show how simCAS tackles the challenge and benefits benchmarking in cis-regulatory interaction inference.

We used synthetic data generated from simCAS fitted by the Buenrostro2018 dataset to illustrate the performance of Cicero (Methods). First, we extended gene regions in chromosome 1 upstream by 50kb and downstream by 50kb to obtain peak hubs. We then filtered the hubs containing less than 50 peaks and the hubs overlapped with others. 14 peak hubs (referred to as hub_1 to hub_14) are retained with the number of peaks ranging from 51 to 92 and the hub length ranging from 337,196 bps to 1,622,989 bps (Fig. 5c). Finally, we randomly selected 40 peaks of each hub to construct interactive peaks, and the remaining peaks are non-interactive samples without modelling correlations.

With the simulated data, we performed Cicero with the window parameter of 200,000 to predict the co-accessibility scores for peak-peak pairs in the simulated data, and the CCANs are constructed by the predicted interactions. Considering the interactive peaks as positive samples and non-interactive peaks as negative samples, we calculated a F1 score for each pair of peak sets between predefined peak hubs and predicted CCANs, and evaluated the performance by the highest F1 score for each peak hub across all the CCANs. As shown in Fig. 5c, Cicero consistently performed well in most peak hubs with different parameters, and achieved the highest F1 score when setting the co-accessibility score cutoff to 0.5, which is reasonable because the possible interactive peaks will be discarded with higher cutoffs and more false positive samples may be predicted with lower cutoffs. With the penalization on the correlations by the distance, Cicero failed to annotate the regions of length significantly exceed the setting window, such as the hub_1 and hub_7 (Fig. 5c). We further visualized the co-accessibility scores predicted by Cicero on the hub_3 region of gene KAZN. As shown in Fig. 5d, with the co-accessibility cutoff of 0.5, most interactive peaks are predicted to be associated with each other, and the interaction is stronger with a closer distance. For the non-interactive peaks, Cicero successfully avoided false positive interactions. With the flexible adaptation of interactive peaks, simCAS is potential to guide computational method benchmarking in cis-regulatory interaction inference.

## 4 Disscusion

We developed a Python package simCAS, an embedding-based simulator of scCAS data, to simulate data from user-defined low-dimensional embeddings. With statistical evaluation and biological evaluation using multiple datasets with different protocols, and with different sizes, dimensions and qualities, we illustrated that simCAS not only preserve cell-wise and peak-wise properties, but also capture biological signals. By testing various analysis methods on simulated data, we also demonstrate the capability of simCAS for benchmarking analysis, suggesting that simCAS has the potential to accelerate development of computational methods for scCAS data analysis.

We also describe several avenues for improving simCAS. First, we can develop a R version of simCAS to offer users the convenience for benchmarking methods programmed in different languages. Second, we can focus on higher-level properties in real scCAS data, such as cell-cell correlation or variance at cell-wise and peak-wise. Finally, due to rapid advances in sequencing technologies, we look forward to extending the framework for single-cell multi-omics data simulation to satisfy the benchmarking for more single-cell related computational methods.

Conflict of Interest: none declared.

