## Supplemental File for "simCAS: an embedding-based method for simulating single-cell chromatin accessibility sequencing data"

[1] Ministry of Education Key Laboratory of Bioinformatics, Bioinformatics Division at the Beijing National Research Center for Information Science and Technology, Center for Synthetic and Systems Biology, Department of Automation, Tsinghua University, Beijing 100084, China

### Supplementary Notes

#### Supplementary Note 1: Adapted simCAS framework with a Bernoulli assumption

Adapted simCAS with a Bernoulli assumption (referred to as simCAS_Bernoulli) is different from the original simCAS (referred to as simCAS_Poisson) in the following steps: estimation for distributions of statistics, parameter and synthetic peak-by-cell matrix generation.

In the step of estimation for distributions of statistics, simCAS_Bernoulli only estimates distributions of library size (the number of aligned reads per cell) and peak summation (the sum of aligned reads per peak), except for cell non-zero proportion (the proportion of non-zero values per cell), which is repeated with library size a Bernoulli assumption. The estimation of library size and peak mean of simCAS_Bernoulli is consistent with simCAS_Poisson.

In the step of parameter matrix correction, simCAS_Bernoulli performs the corrections by estimated peak mean and library size. For simplicity, here we used the symbols with the same meaning in simCAS_Poission. Suppose $n_{cell}$, $n_{peak}$ and $n'_{cell}$ denote number of simulated cells, number of simulated peaks and number of referenced cells in real data, respectively. After the low-dimensional embeddings generation and activation transformation as in simCAS_Poisson, we obtain the activated parameter matrix $\widehat{\boldsymbol{\Lambda}} \in \mathbb{R}^{n_{peak} \times n_{cell}}$. The library size correction is performed as follows:

(1) Multiply the CEV $\boldsymbol{l}$ and the CEM $\boldsymbol{C}$, and obtain $\hat{\boldsymbol{l}} \in \mathbb{R}^{1 \times n_{cell}}$. Each value of $\hat{\boldsymbol{l}}$ represents the weight of potential library size of each column vector (represents each cell) in $\widehat{\boldsymbol{\Lambda}}$.

(2) Randomly sample $n_{cell}$ values from the GMM distribution fitted with real library sizes to form the synthetic library size set $L \in \mathbb{R}^{n_{cell}}$. Then assign the values of $L$ to each column vector in $\widehat{\boldsymbol{\Lambda}}$ with the same order of the potential library size weights. Denote the value of $L$ assigned to $j$th cell of $\widehat{\boldsymbol{\Lambda}}$ as $l_j$.

(3) For each column in $\widehat{\boldsymbol{\Lambda}}$ we perform a uniform correction. For the $j$th column vector $\hat{\boldsymbol{\lambda}}_{.,j}$ in $\widehat{\boldsymbol{\Lambda}}$, sort it with an ascending order to get $\hat{\boldsymbol{\lambda}}_{.,j}^{sort}$. Then search from the largest value to the lowest value of $\hat{\boldsymbol{\lambda}}_{.,j}^{sort}$ to find an index $I_j$ and a factor $k_j$ to satisfy the equations:

$$\begin{cases} (\Sigma_{i=1}^{I_j}\hat{\lambda}_{i,j}^{sort}k_j) + (n_{peak} - I_j) = l_j \\ \hat{\lambda}_{I_j,j}^{sort}k_j < 1 \\ \hat{\lambda}_{I_j+1,j}^{sort}k_j \geq 1 \end{cases},$$

where $\hat{\lambda}_{i,j}^{sort}$ is the $i$th value of $\hat{\boldsymbol{\lambda}}_{\cdot,j}^{sort}$.

(4) Then multiply $\hat{\boldsymbol{\lambda}}_{\cdot,j}$ with $k_j$ and set the values exceeding 1 to 1 to get the corrected parameters.

(5) Conduct the uniform correction for each column in $\widehat{\boldsymbol{\Lambda}}$ and acquire the library size corrected parameter matrix $\check{\boldsymbol{\Lambda}}$.

The peak mean correction is conducted with a similar operation:

(1) Randomly sample $n_{peak}$ values from the estimated discrete distribution fitted with real peak summations to form the synthetic peak summation set $P \in \mathbb{R}^{n_{peak}}$. Then assign the values of $P$ to each row vector (represents each peak) in $\check{\boldsymbol{\Lambda}}$ with the same order of the peak-wise summations in $\widehat{\boldsymbol{\Lambda}}$. Denote the value of $P$ assigned to $i$th peak of $\widehat{\boldsymbol{\Lambda}}$ as $p_j$

(2) For each row in $\check{\boldsymbol{\Lambda}}$ we perform a uniform correction. For the $i$th row vector $\check{\boldsymbol{\lambda}}_{i,\cdot}$ in $\check{\boldsymbol{\Lambda}}$, sort it with an ascending order to get $\check{\boldsymbol{\lambda}}_{i,\cdot}^{sort}$. Then search from the largest value to the lowest value of $\check{\boldsymbol{\lambda}}_{i,\cdot}^{sort}$ to find an index $I_i$ and a factor $k_i$ to satisfy the equations:

$$\begin{cases} \left(\Sigma_{j=1}^{I_i}\check{\lambda}_{i,j}^{sort}k_i\right) + (n_{cell} - I_i) = n_{cell}\dfrac{p_j}{n'_{cell}} \\ \check{\lambda}_{i,I_i}^{sort}k_i < 1 \\ \check{\lambda}_{i,I_i+1}^{sort}k_i \geq 1 \end{cases},$$

where $\check{\lambda}_{i,j}^{sort}$ is the $j$th value of $\check{\boldsymbol{\lambda}}_{i,\cdot}^{sort}$.

(3) Then multiply $\check{\boldsymbol{\lambda}}_{i,\cdot}$ with $k_i$ and set the values exceeding 1 to 1 to get the corrected parameters.

(4) Conduct the uniform correction for each row in $\check{\boldsymbol{\Lambda}}$ and acquire the peak mean corrected parameter matrix $\boldsymbol{\Lambda}$.

In the step of synthetic peak-by-cell matrix generation, we replace the Poisson distribution with the Bernoulli distribution, of which the probability parameter is the corresponding parameter in $\boldsymbol{\Lambda}$.

**Supplementary Note 2: Alternative distributions for modeling peak summation**

We provide five discrete distributions for modeling peak summation as alternatives, namely Log-variantB (another variant of Logarithmic distribution different with Log-variant), NB (Negative Binomial), ZINB (Zero-Inflated Negative Binomial), NB-variant (a variant of Negative Binomial distribution) and ZIP distributions. For simplicity of notation, we assume $x$ as a random variable with no practical meaning in the following discussion.

The probability mass function (PMF) of the Log-vairantB distribution is:

$$
\begin{aligned}
&f_{\mathrm{Log-vairantB}}(x; p, \pi)\\
&= \pi_0 \delta_0(x) + (1 - \pi_0) f_{\mathrm{Logarithmic}}(x; p)\\
&= \pi_0 \delta_0(x) + (1 - \pi_0) \frac{-1}{\ln(1-p)} \frac{p^x}{x}
\end{aligned}
$$

where $\delta_0(\cdot)$ indicates the point mass at zero, and $\pi_0$ is the corresponding probability. $p$ is a parameter ranging from 0 to 1 in Logarithmic distribution.

The PMF of the NB distribution is:

$$f_{\mathrm{NB}}(x; r, p_0) = \frac{(x + r - 1)!}{(r-1)!\, x!} p_0^x (1 - p_0)^x$$

where $r$ denotes the number of successes, and $p_0$ denotes the probability of success on each trial.

The PMF of the ZINB distribution is:

$$f_{\mathrm{ZINB}}(x; \pi_0, r, p_0) = \pi_0 \delta_0(x) + (1 - \pi_0) f_{\mathrm{NB}}(x; r, p_0)$$

The PMF of the NB-variant distribution is:

$$f_{\mathrm{NB-variant}}(x; \pi_0, r, p_0) = \pi_0 \delta_0(x) + (1 - \pi_0) f_{\mathrm{NB}}(x - 1; r, p_0)$$

The PMF of the ZIP distribution is:

$$f_{\mathrm{ZIP}}(x; \pi_0, r, p_0) = \pi_0 \delta_0(x) + (1 - \pi_0) f_{\mathrm{Poisson}}(x; \lambda) = \pi_0 \delta_0(x) + (1 - \pi_0) \frac{\lambda^x e^{-\lambda}}{x!}$$

where $\lambda$ denotes the mean parameter of Poisson distribution.

#### Supplementary Note 3: Correction of peak summation and cell sparsity

After correction of library size, simCAS obtains the corrected parameter matrix $\breve{\boldsymbol{\Lambda}}$. simCAS then performs the peak summation correction as follows:

(1) Randomly sample $n_{peak}$ values from the Log-variant distribution fitted with real peak summation to form the synthetic peak summation set $P \in \mathbb{R}^{n_{peak}}$.

(2) Sum the elements of each row vector in $\breve{\boldsymbol{\Lambda}}$, and obtain $\breve{\boldsymbol{p}} \in \mathbb{R}^{n_{peak}\times 1}$

(3) Sort the elements of $\breve{\boldsymbol{p}}$ and values in $P$ to obtain the sorting indices.

(4) Obtain the vector $\breve{\boldsymbol{p}}' \in \mathbb{R}^{n_{peak}\times 1}$ of the peak summations of synthetic cells by replacing the elements in $\breve{\boldsymbol{p}}$ with the sampled values from $P$ one by one according to the sorting indices.

(5) For each row vector (represents each peak) in $\breve{\boldsymbol{\Lambda}}$, divide each element by the sum of the vector, and then multiply all elements by the corresponding element (represents the associated cell) in $\breve{\boldsymbol{p}}'$. Then obtain the corrected parameter matrix $\acute{\boldsymbol{\Lambda}}$.

After correction of peak summation, simCAS obtains the corrected parameter matrix $\acute{\boldsymbol{\Lambda}}$. simCAS finally performs the cell sparsity correction as follows:

(1) Randomly sample $n_{cell}$ values from the GMM distribution fitted with real cell sparsity to form the synthetic cell sparsity set $S \in \mathbb{R}^{n_{cell}}$.

(2) For the $j$th synthetic cell, randomly select a value in $S$ without replacement, called $s_j$, as the sparsity of the cell.

(3) For the $j$th synthetic cell, obtain $k_j$ and $\pi_j$ by solving two nonlinear equations as follows:

$$\begin{cases} \sum_{m=1}^{n_{cell}} (1-\pi_j)(\acute{\lambda}_{m,j} k_j) = \sum_{m=1}^{n_{cell}} \acute{\lambda}_{m,j} \\ n_{peak}\pi_j + (1-\pi_j) \sum_{m=1}^{n_{cell}} e^{-\acute{\lambda}_{m,j} k_j} = n_{peak}(1-s_j) \end{cases}$$

where $k_j$ and $\pi_j$ denote a scaling factor and a zero-setting probability of the $j$th synthetic cell, respectively, and $\acute{\lambda}_{m,j}$ is the element in $\acute{\mathbf{\Lambda}}$ corresponding to the $m$th peak and the $j$th cell.

(4) For the $j$th synthetic cell, randomly set elements of the corresponding column vector to zero with the probability $\pi_j$, and then multiple the remaining non-zero elements by $k_j$.

(5) For each cell, performing the processing from (2) to (4), obtain the final mean parameter matrix $\mathbf{\Lambda}$.

#### Supplementary Note 4: Details of the optional steps in simCAS

simCAS could generates multi-batch data by incorporating the batch effects in the discrete simulation mode. For simulating data with the technical batch effect, we add the Gaussian noise on the corrected parameter matrix $\mathbf{\Lambda}$:

$$\mathbf{\Lambda}' = \mathbf{\Lambda} + \varepsilon_1 ,$$

where $\varepsilon_1$ is Gaussian noise sampled from $\mathcal{N}(\mu_t, \sigma_t^2)$. $\varepsilon_1$ can be deemed as technical variations when sequencing reads and results in an evident variation in library size values, and $\mu_t$ and $\sigma_t$ control the variance of batches and degree of technical noise, respectively. The common CEM of data generated from $\mathbf{\Lambda}'$ and $\mathbf{\Lambda}$ guarantees the ground truth of same cells from different batches. With different levels of Gaussian noise added on $\mathbf{\Lambda}$, data with multiple technical batch effects can be generated. Note that the technical variance may be different in the cell types with different library sizes, we add Gaussian noise proportion to the average cell-wise summations of $\mathbf{\Lambda}$ in different cell populations.

The biological batch effect can be directly modeled by adding Gaussian noise to the PEM:

$$\boldsymbol{P}' = \boldsymbol{P} + \varepsilon_2 ,$$

where $\varepsilon_2$ is Gaussian noise sampled from $\mathcal{N}(\mu_b, \sigma_b^2)$. Then synthetic data with different batches is generated with a common CEM, which provides the ground truth of cells with same population.

The interactive peaks are constructed by assigning similar vectors in the PEM generation step. First, we pick up high chromatin accessibility regions to define hubs of interactive peaks, and then the correlation is added on the peaks of each hub. In a defined peak hub, randomly select some peaks as the interactive peaks and remaining peaks are non-interactive peaks. For interactive peaks in the hub, we first generate a general peak embedding vector $\widehat{\boldsymbol{p}} \in \mathrm{R}^{1\times n_{embed}}$, then the embedding vector of each interactive peak is generated by:

$$\boldsymbol{p}_{j'} = \widehat{\boldsymbol{p}} + \varepsilon_3 ,$$

where $\varepsilon_3$ is sampled from $\mathcal{N}(0, \sigma_p^2)$ and $\sigma_p$ is a parameter to control the degree of co-accessibility among interactive peaks.

**Supplementary Note 5: Evaluation metrics**

Suppose $R$ is the sorted real statistic values $S$ is the sorted simulated statistic values, MAD, MAE and RMSE are calculated as following equations:

$$MAD(R,S) = median(|R - S|)$$

$$MAE(R,S) = mean(|R - S|)$$

$$RMSE(R,S) = \sqrt{mean((R - S)^2)}$$

PCC calculates linear correlation between $R$ and $S$:

$$PCC(R,S) = \frac{\mathrm{cov}(R,S)}{\sigma_R \sigma_S},$$

where cov is the variance and $\sigma$ is the standard deviation.

JSD is a measurement of the similarity between two probability distributions. For the reason that integral of continuous variable is incapable be calculated directly, JSD is merely calculated to compare discrete statistic peak mean:

$$JSD(R,S) = \frac{1}{2}D(R||M) + \frac{1}{2}D(S||M)$$

$$M = \frac{1}{2}(R + S)$$

$$D(P||Q) = \Sigma_x P(x)\log\left(\frac{P(x)}{Q(x)}\right),$$

where $D$ is Kullback–Leibler divergence, $P$ and $Q$ are two probability distributions.

KS statistic quantifies the distance between the empirical distribution of two samples, which is derived from a nonparametric test of the equality of two continuous samples named KS test. We apply it on continuous statistics such as the log-transformed library size and the cell sparsity.

Median integration local inverse Simpson's index (miLISI) is a score first proposed to evaluate the integration of data with different batches. Gaussian kernel-based distributions of neighborhoods of the mixing batches are built in low-dimensional embedding space. iLISI is then computed for each neighborhood:

$$\mathrm{iLISI} = \frac{1}{\Sigma_{b=1}^{B} p(b)},$$

where $p(b)$ refers to the probability that two sampling neighbors are from the same batch $b$

and $B$ is the number of batches. By considering original data and synthetic data as different batches, this score can directly be adapted to quantify the similarity between synthetic cells and real cells. This score ranges from 1 to 2, and a larger value indicates a greater similarity. The closer miLISI is to 2, the local neighborhood has more equal synthetic and real cells. We calculate miLISI value on the 2D UMAP embedding space containing real and synthetic cells using the R package LISI.

Denoting ground truth with $gt$ and clustering labels with $pred$. Homo is calculated using:

$$Homo(gt, pred) = 1 - \frac{H(gt|pred)}{H(gt)},$$

where $H$ is the entropy and $H(gt|pred)$ is the conditional entropy of ground truth clusters given the unsupervised predictions.

AMI is calculated using:

$$AMI(gt, pred) = \frac{MI - E[MI]}{mean\big(H(gt), H(pred)\big) - E[MI]},$$

where $MI$ is the mutual information and $E(\cdot)$ is the expectation function.

ARI is calculated using:

$$ARI(gt, pred) = \frac{RI - E[RI]}{\max(RI) - E[RI]},$$

where Rand index (RI) is a similarity measurement between ground truth labels and predicted labels.

F1 score is calculated as follows:

$$\mathrm{F1} = \frac{TP}{TP + \frac{1}{2}(FP + FN)},$$

where $TP$, $FP$, $TN$, $FN$ represent true positive, false positive, true negative and false negative, respectively.

#### Supplementary Figures

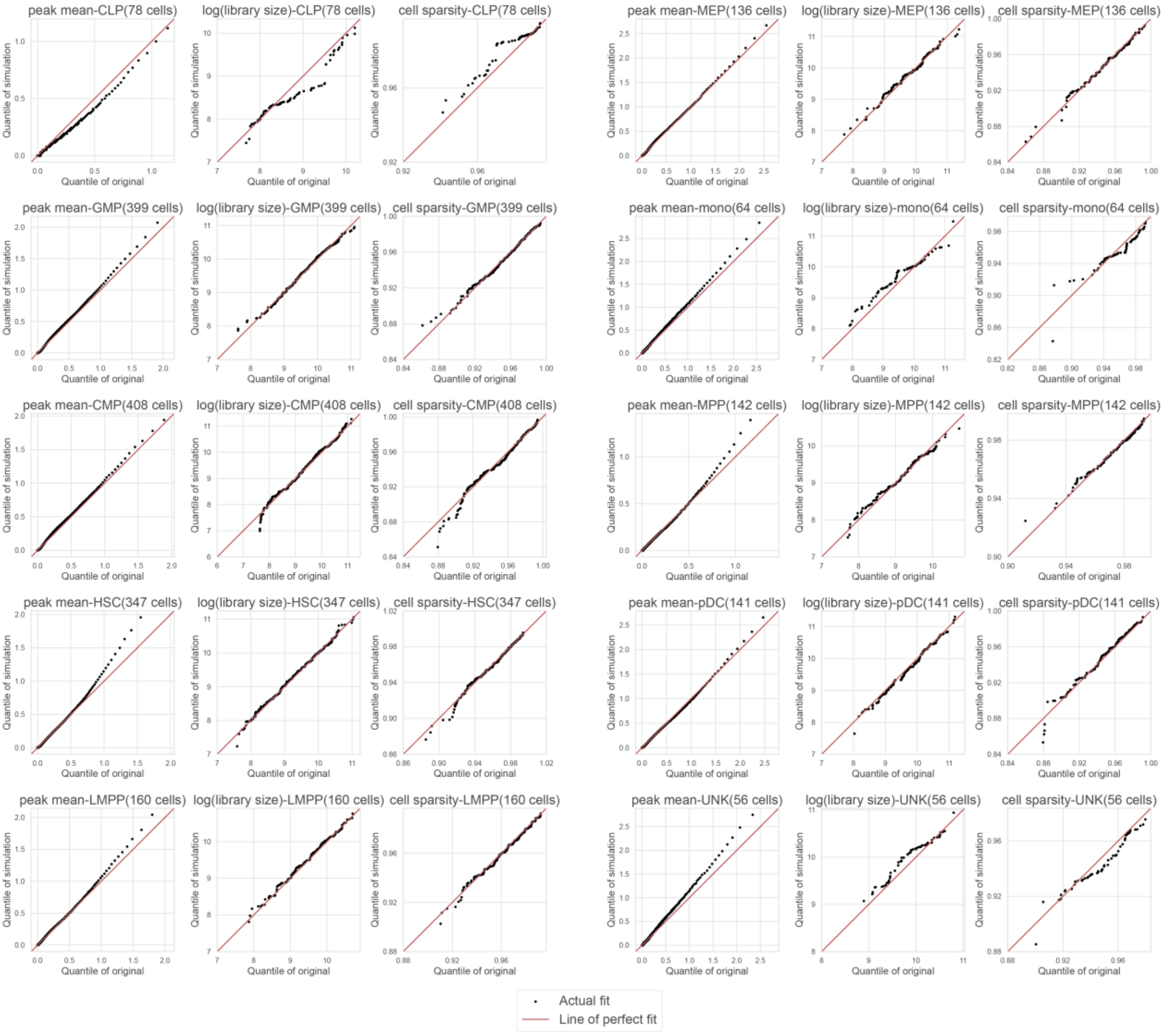

**Supplementary Fig. 1.** The comparison of the distributions of peak mean, library size and cell sparsity between real data and synthetic data for 10 cell types in Buenrostro2018 dataset.

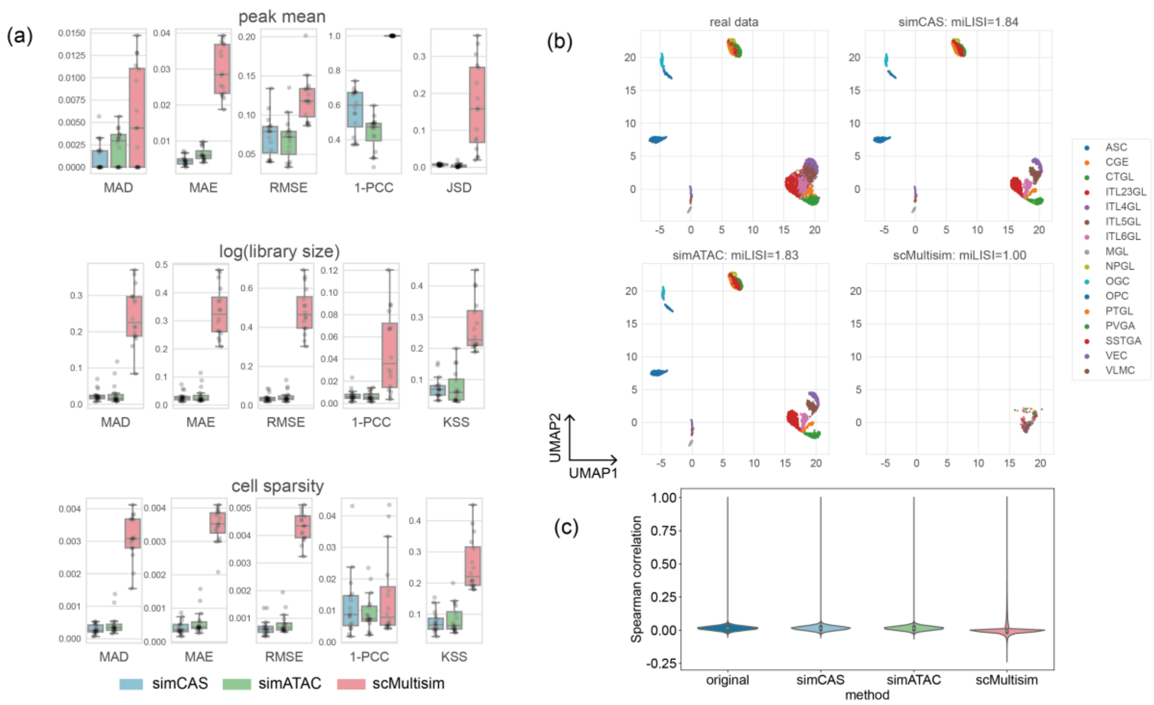

**Supplementary Fig. 2.** Comparisons of synthetic data and real data in Li2021 dataset. (b) Comparison results of three statistics between synthetic cell types and real cell types measured by six metrics. (c) UMAP visualization of the synthetic datasets and real dataset. (d) Spearman correlation coefficients of synthetic and real datasets on the pairs of top 2000 highly variable peaks selected in real data.

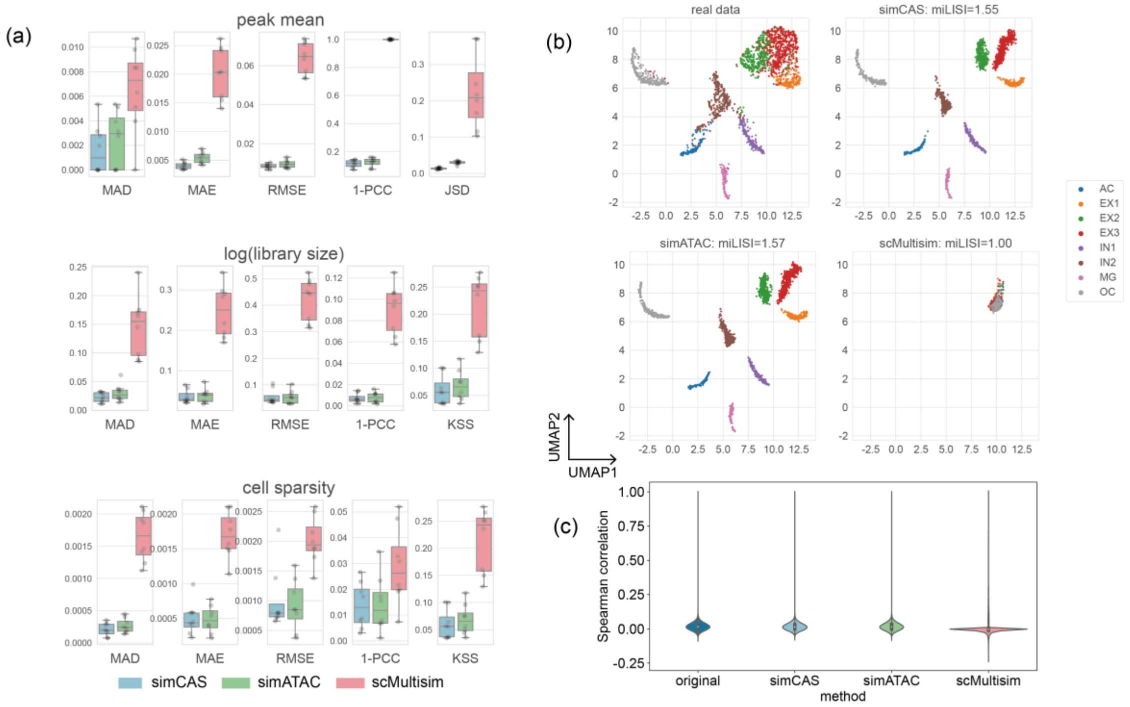

**Supplementary Fig. 3.** Comparisons of synthetic data and real data in Pressl2018 dataset. (b) Comparison results of three statistics between synthetic cell types and real cell types measured by six metrics. (c) UMAP visualization of the synthetic datasets and real dataset. (d) Spearman correlation coefficients of synthetic and real datasets on the pairs of top 2000 highly variable peaks selected in real data.

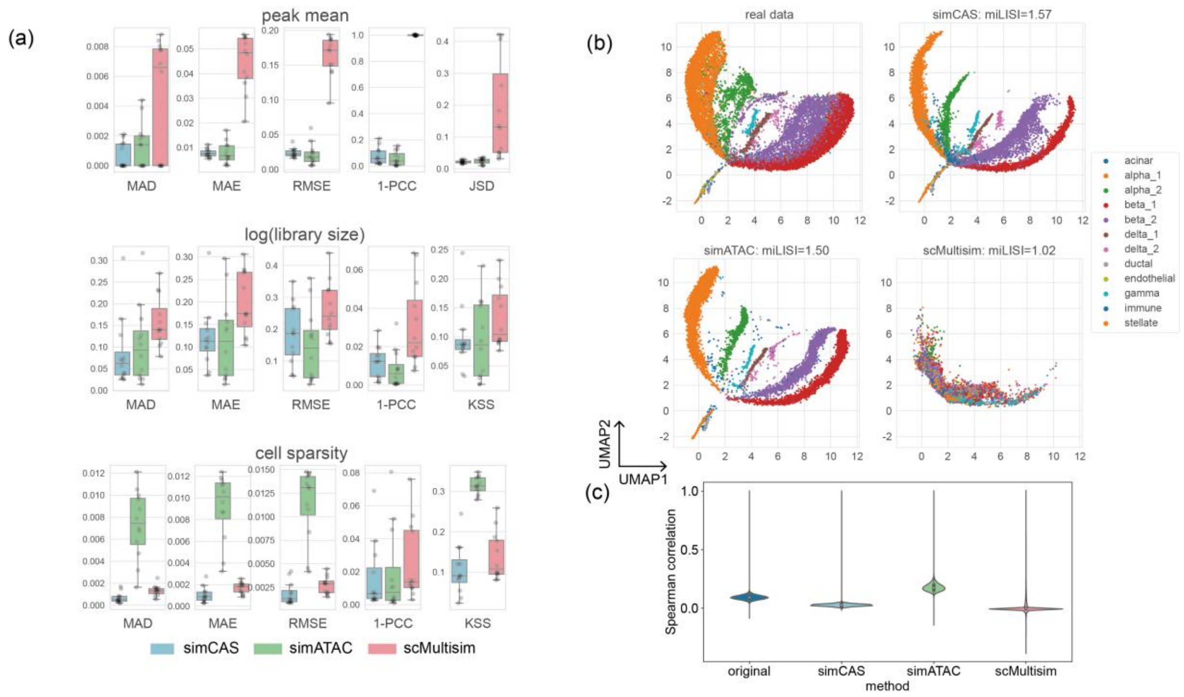

**Supplementary Fig. 4.** Comparisons of synthetic data and real data in Chiou2021 dataset. (b) Comparison results of three statistics between synthetic cell types and real cell types measured by six metrics. (c) UMAP visualization of the synthetic datasets and real dataset. (d) Spearman correlation coefficients of synthetic and real datasets on the pairs of top 2000 highly variable peaks selected in real data.

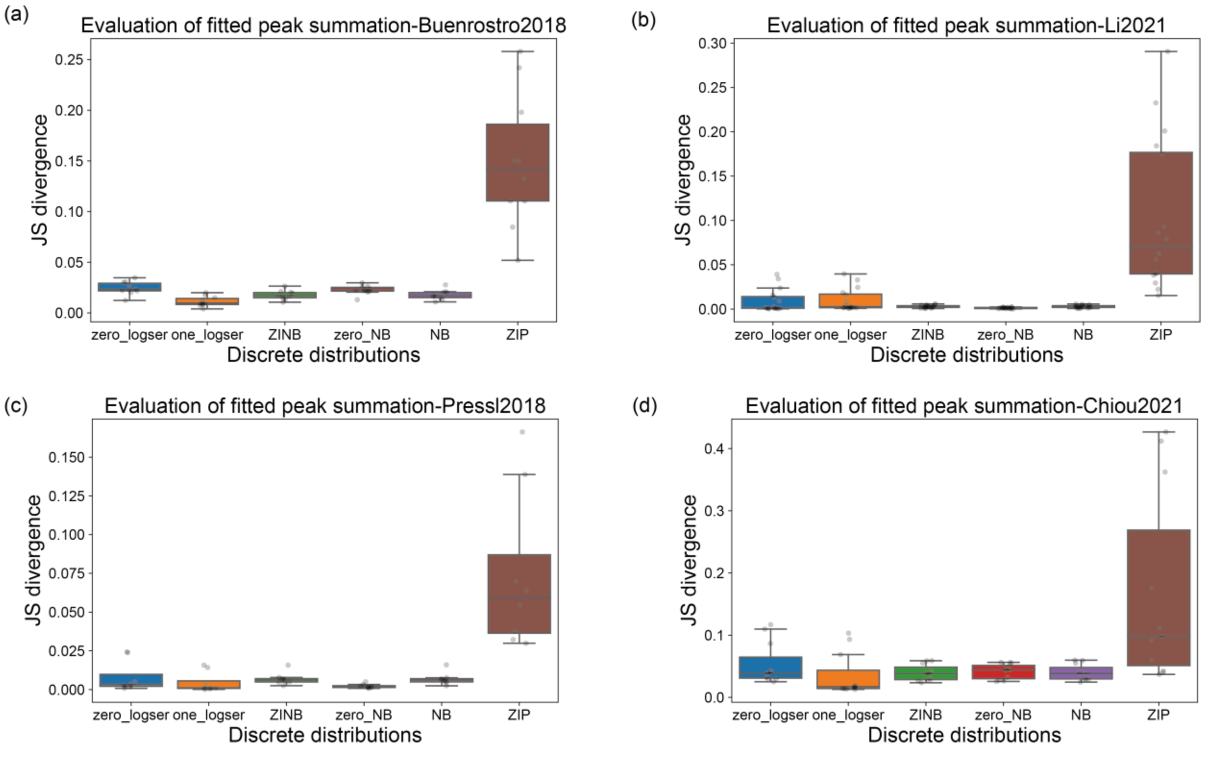

**Supplementary Fig. 5.** Estimation performance of six discrete distributions on fitting the peak summation of real datasets. The estimation performance is measured by JS divergence. Four real datasets are utilized in the evaluation: (a) Buenrostro2018 dataset, (b) Li2021 dataset, (c) Pressl2018 dataset, and (d) Chiou2021 dataset.

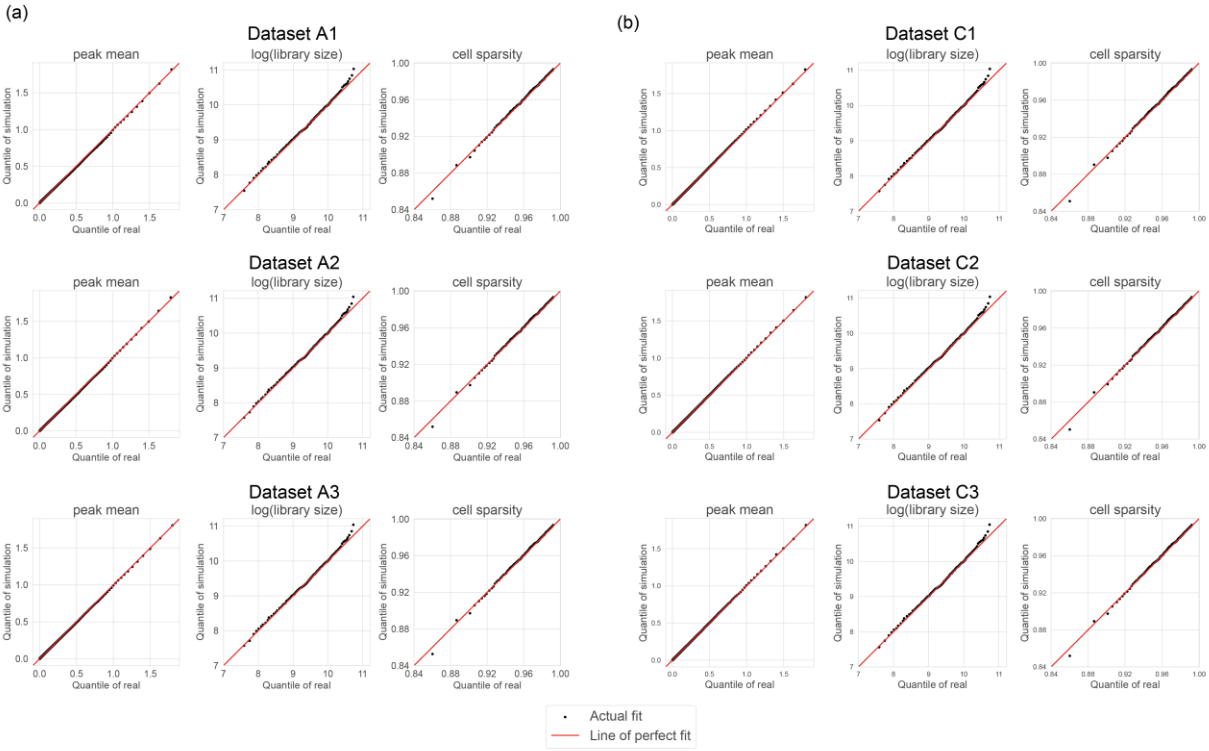

**Supplementary Fig. 6.** QQ-plots of statistics' comparison between real data and synthetic data. (a) Comparison for datasets A1-A3. (b) Comparison for datasets C1-C3.

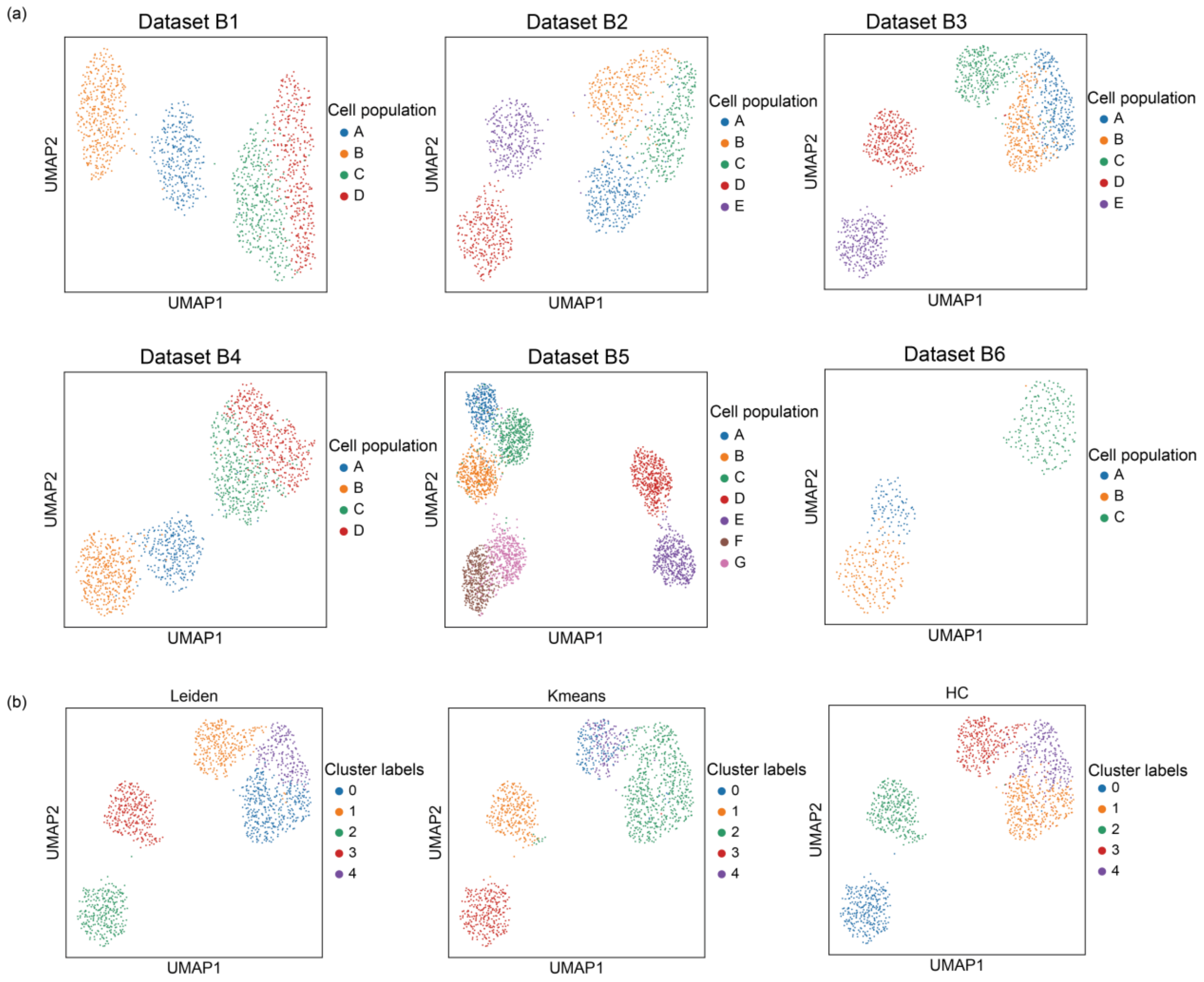

**Supplementary Fig. 7.** UMAP visualization of the simulated datasets for cell clustering benchmarking and the clustering result. (a) UMAP visualization of datasets B1-B6, colored by the predefined cell populations. (b) An illustration of the clustering result of Leiden clustering, K-means clustering and hierarchical clustering applied on the dataset B3.

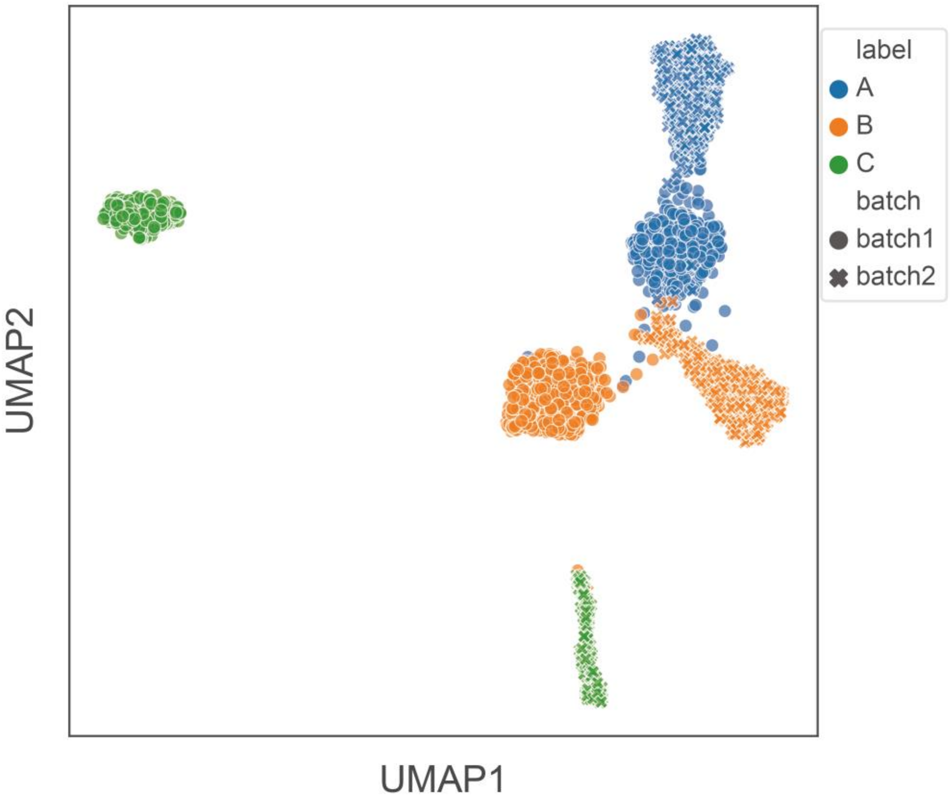

**Supplementary Fig. 8.** UMAP visualization of a synthetic dataset with simulated technical batch effect. Data of batch1 is generated with populations of 3 (the number of cells in cell population A, B and C is set to 600, 600 and 300, respectively) and a unit diagonal matrix as the covariance matrix. Denote $\mathbf{\Lambda}_i$ is part of $\mathbf{\Lambda}$ specifically for $i$th cell population, add Gaussian noise on $\mathbf{\Lambda}_i$ proportional to the average cell-wise summations of $\mathbf{\Lambda}_i$ to generate data batch2 with technical batch effect compared to data batch1. In this experiment the Gaussian noise is added with mean of 1.88, 0.66 and 0.50 for population A, B and C, respectively, and a fixed standard deviation of 0.5. Then the matrices of data batch1 and data batch2 are concatenated to acquire the data with technical batch effect.
